## Supplemental Figures for "Lethality of SARS-CoV-2 infection in K18 human angiotensin converting enzyme 2 transgenic mice"

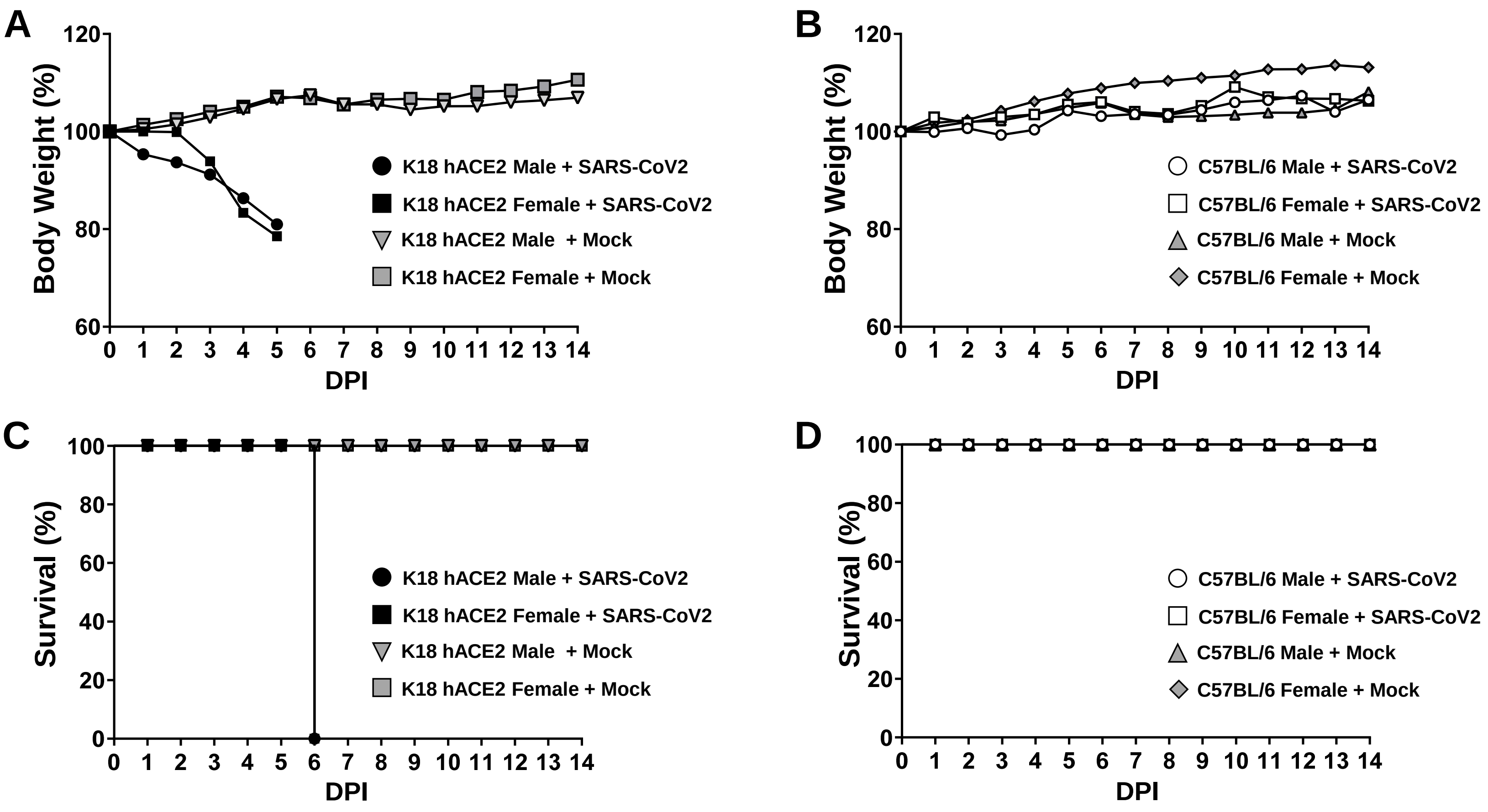


**Figure S1. Infection of K18 hACE2 transgenic and WT C57BL/6 mice with SARS-CoV-2.** Female and male K18 hACE2 transgenic and WT C57BL/6 mice were mock-infected (N=3/group) or infected (N=4/group) i.n. with 1 x $10^5$ PFU of SARS-CoV-2. Body weight (**A, B**) and survival (**C, D**) were evaluated at the indicated DPI. Mice that loss more than 25% of their initial body weight were humanely euthanized. Error bars represent standard deviations (SD) of the mean for each group of mice. DPI: Days post-infection.

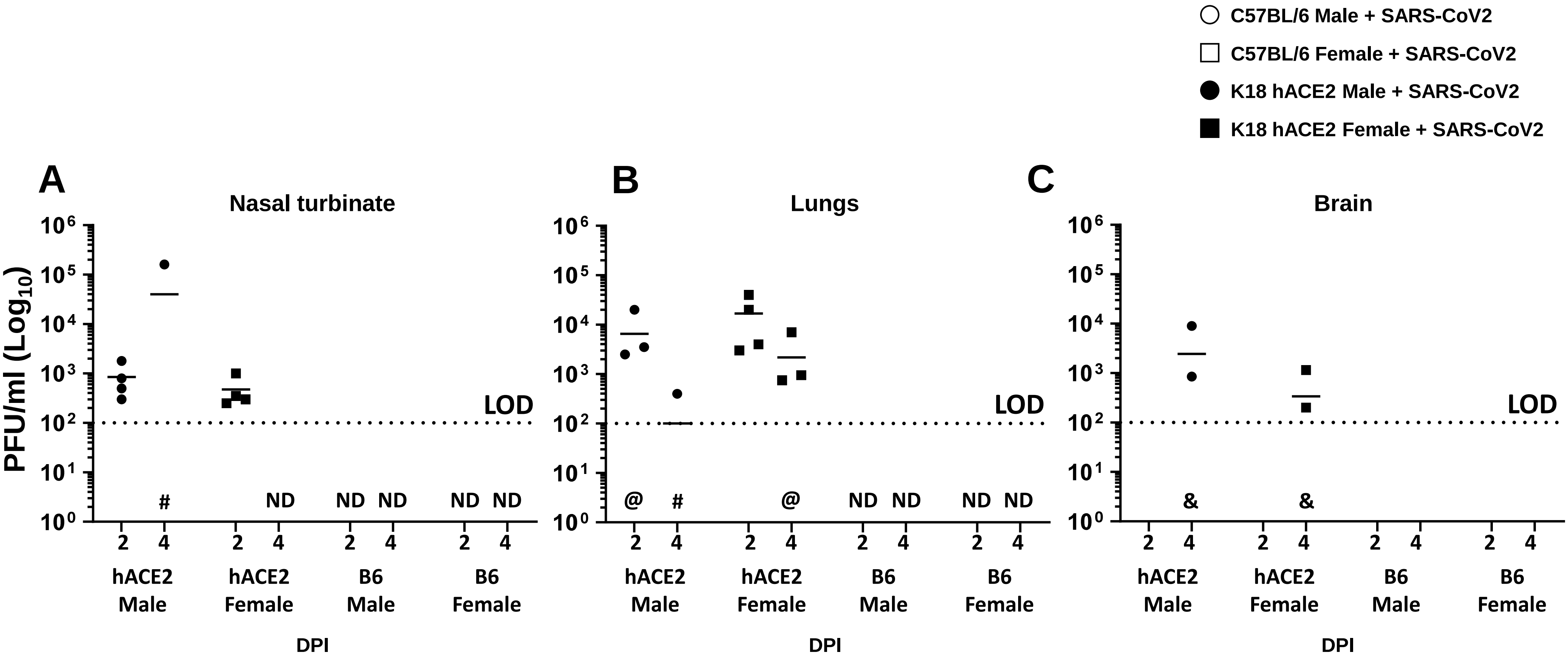


**Figure S2. Viral loads in male and female K18 hACE2 transgenic and WT C57BL/6 mice infected with SARS-CoV-2.** K18 hACE2 transgenic and WT C57BL/6 male (N=4/group) and female (N=4/group) mice were infected as in Figure S1 and sacrificed at 2- and 4-DPI and viral titers in different organs (nasal turbinate, trachea, lung, brain, heart, kidney, liver, spleen, small intestine, and large intestine) were determined by plaque assay (PFU/ml). Only data from virus containing organs and/or tissue samples are shown: nasal turbinate (**A**), lungs (**B**) and brain (**C**). Symbols represent data from individual mouse, and bars the geometric means of viral titers. @, virus not detected in one mouse; &, virus not detected in two mice; #, virus not detected in three mice; ND, not detected. Dotted black lines indicate the limit of detection ($10^2$ PFU/ml). DPI: Days post-infection.

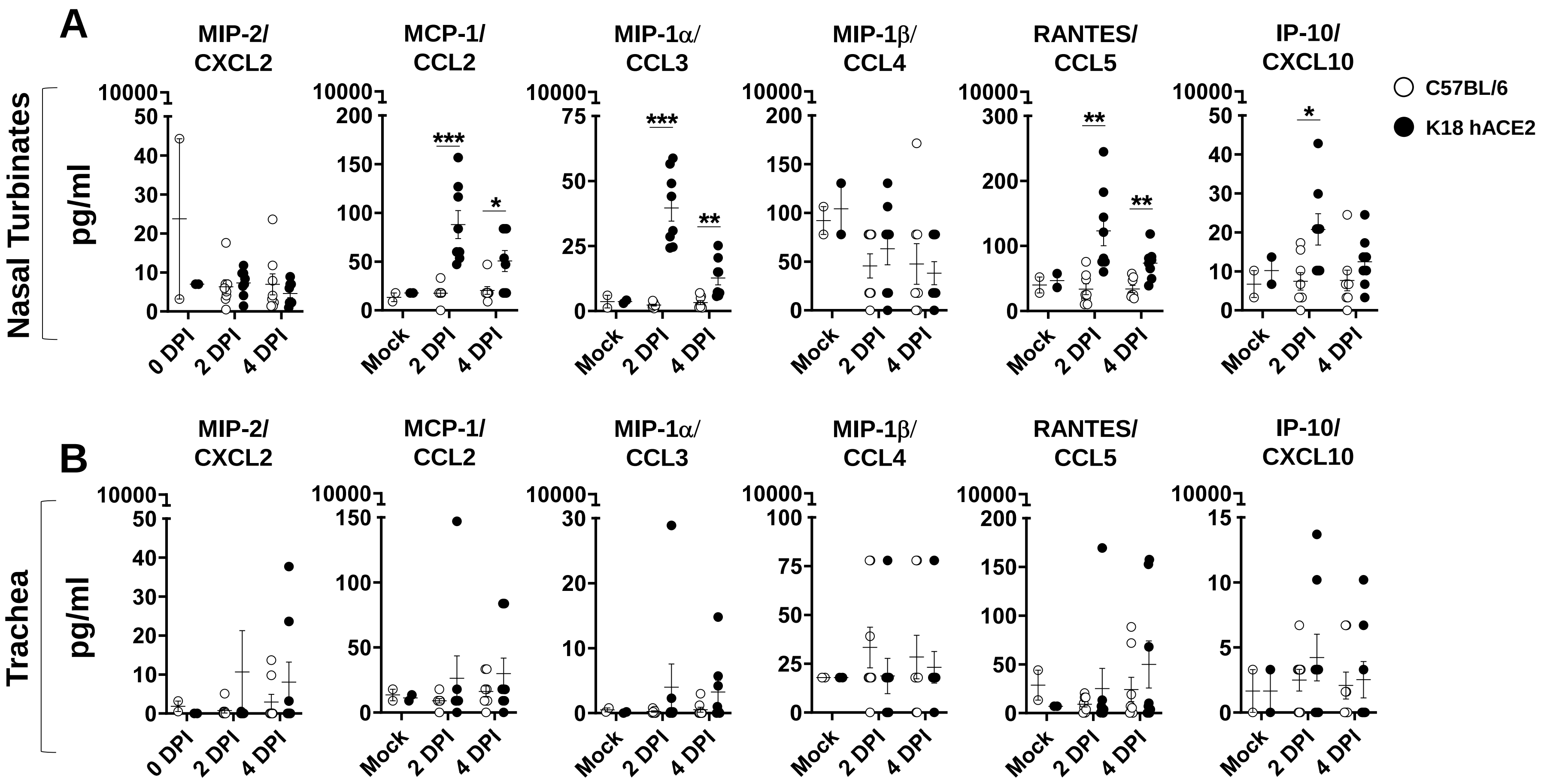


**Figure S3. Chemokine profile in selected tissues from SARS-CoV-2 infected K18 hACE2 transgenic mice.** (**A**) Nasal turbinates and (**B**) Trachea. Student's *t*-test C57BL/6 *vs.* K18 hACE2 **p*< 0.05; ***p*<0.005; ****p*<0.0005; 2-WAY ANOVA C57BL/6 or K18 hACE2 transgenic mice over time, §p< 0.05; §§p<0.005; §§§p<0.0005, N= 8 (per time-point studied, except mock N=2). DPI: Days post-infection.

### Nasal Turbinates

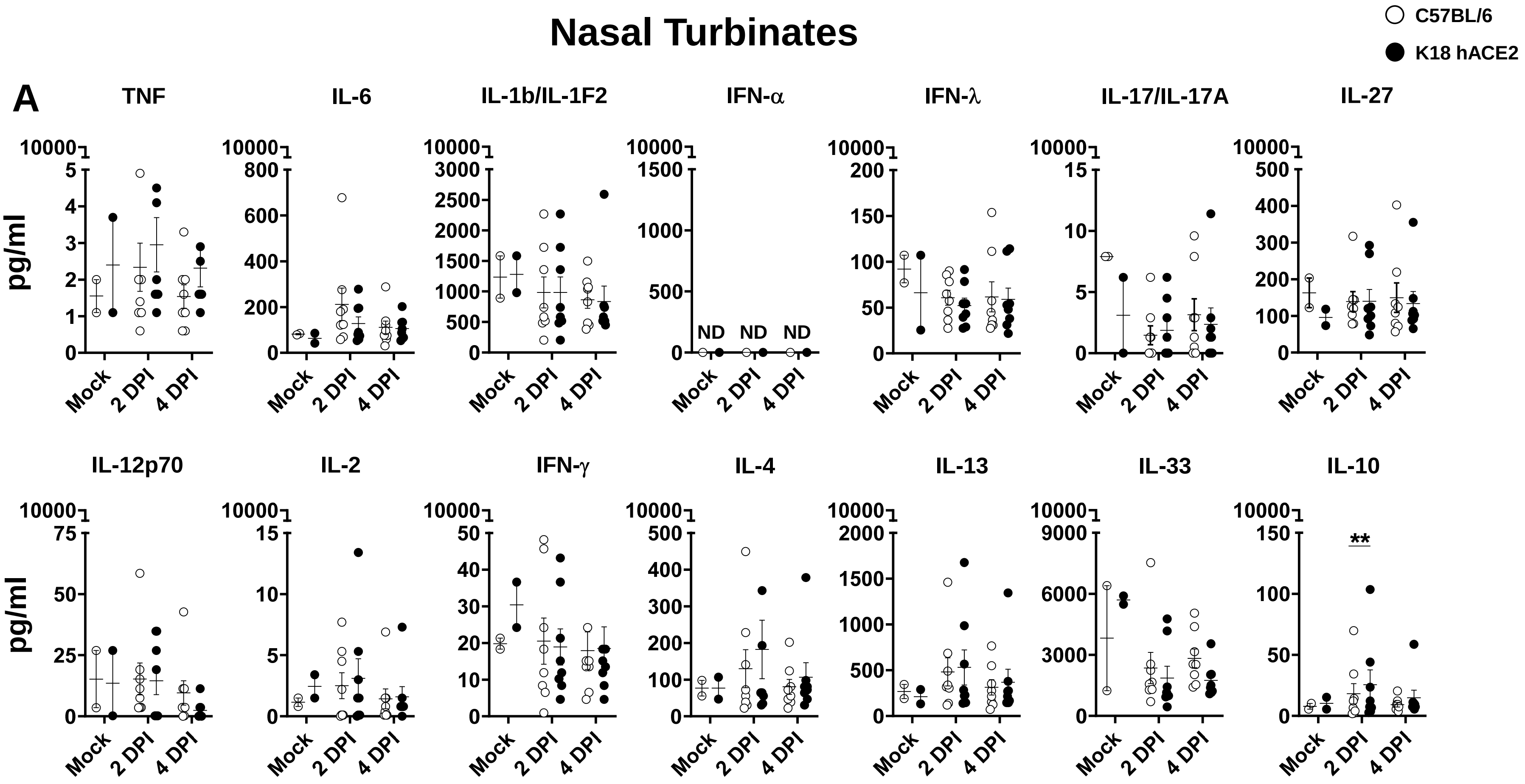


**Figure S4. Cytokine profile in selected tissues from SARS-CoV-2 infected K18 hACE2 transgenic mice.** (**A**) Nasal turbinates and (**B**) Trachea. Student's t-test C57BL/6 *vs.* K18 transgenic hACE2 **p*< 0.05; ***p*<0.005; ****p*<0.0005; 2-WAY ANOVA C57BL/6 or K18 hACE2 transgenic mice over time, §p< 0.05; §§p<0.005; §§§p<0.0005, N= 8 (per time-point studied, except mock N=2). DPI: Days post-infection.

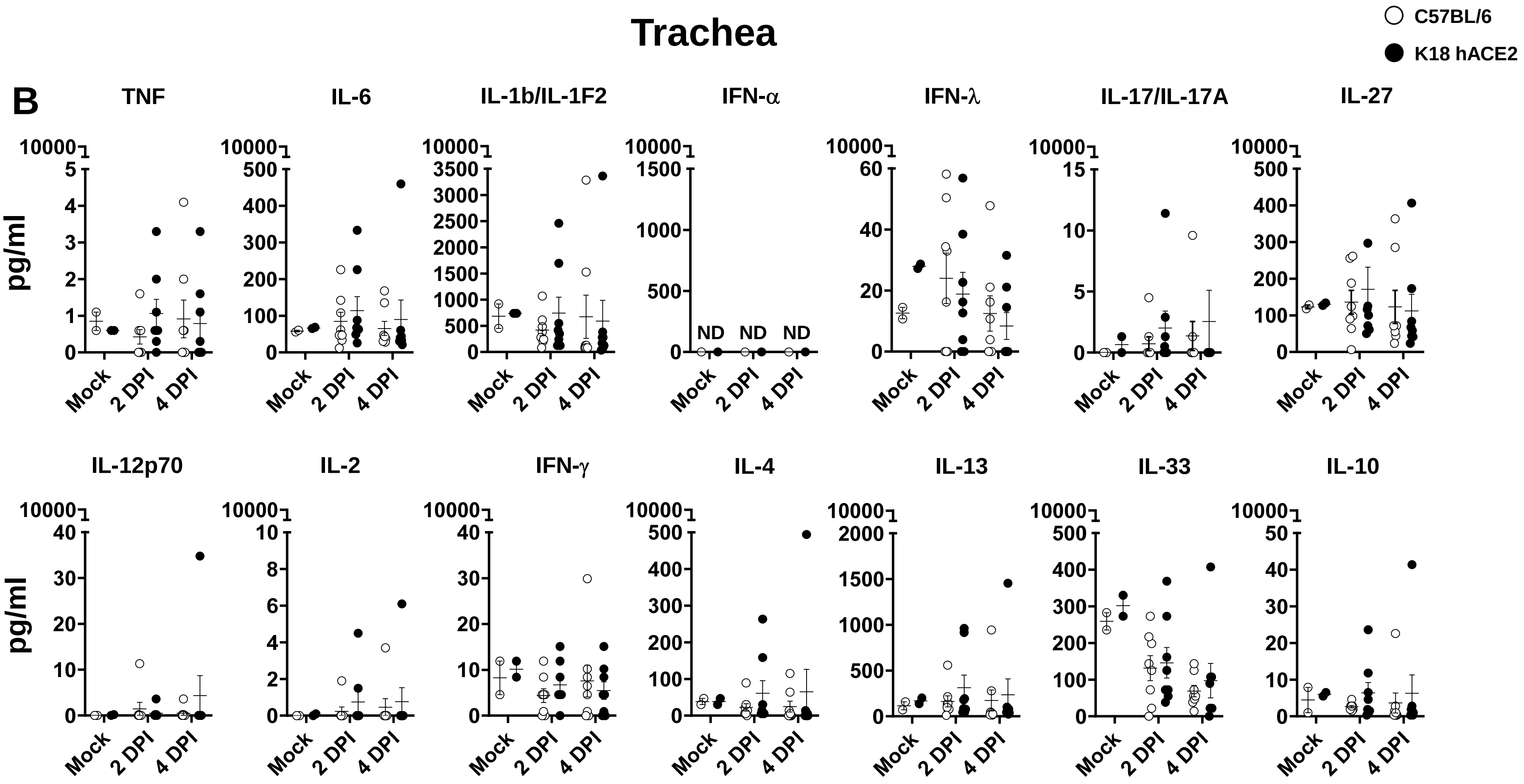


Figure S4. Continuation.

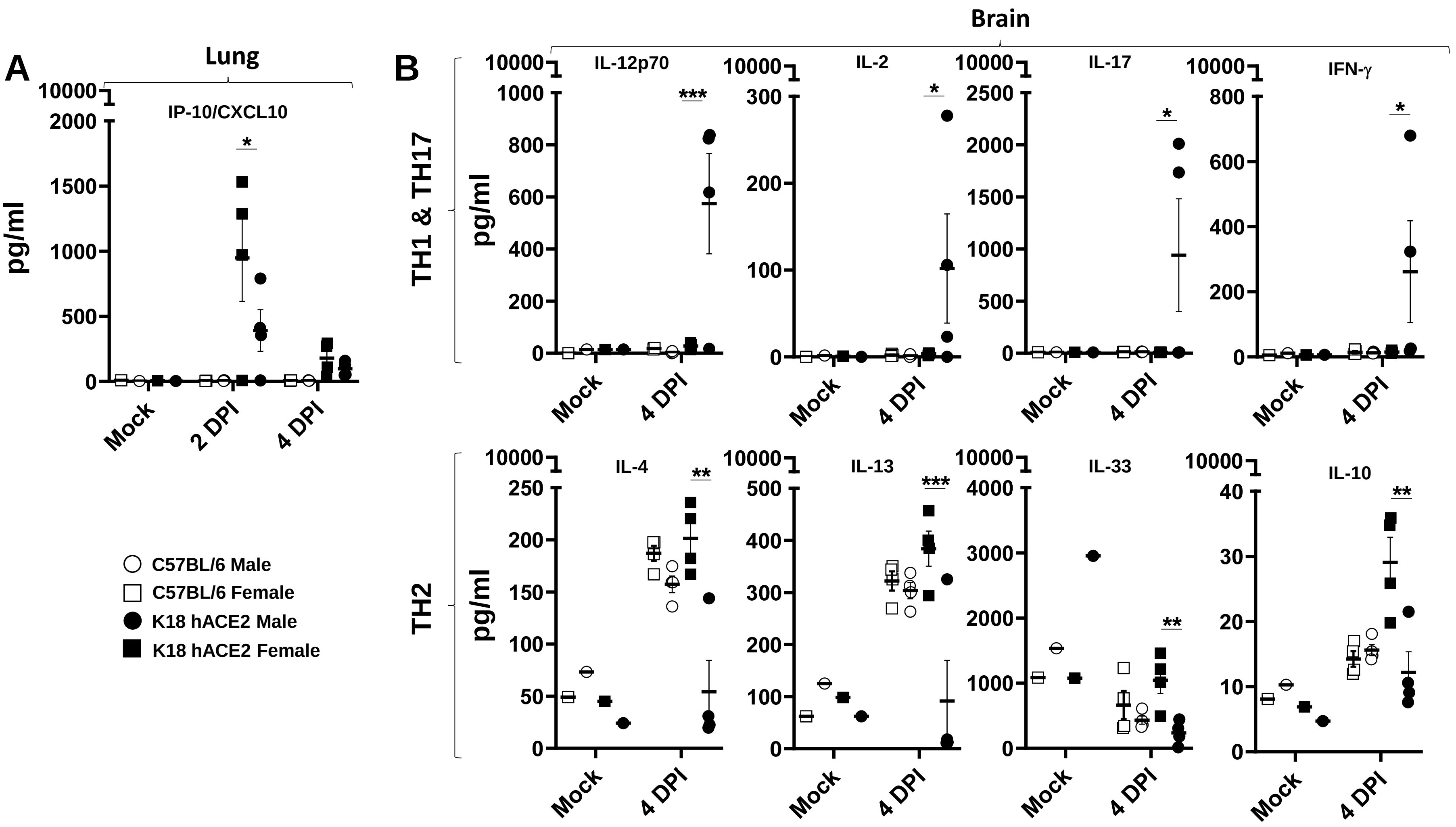


**Figure S5. Chemokine and cytokine profile in selected tissues from SARS-CoV-2 infected K18 hACE2 transgenic mice by sex.** Chemokines and cytokine differences in the (**A**) lung and (**B**) brain in K18 hACE2 transgenic and WT C57BL/6 male and female mice. Student's t-test C57BL/6 *vs.* K18 hACE2 *$p$< 0.05; **$p$<0.005; ***$p$<0.0005; 2-WAY ANOVA C57BL/6 or K18 hACE2 transgenic mice over time, §p< 0.05; §§p<0.005; §§§p<0.0005, N= 4/group/sex (per time-point studied, except mock N=2). DPI: Days post-infection.

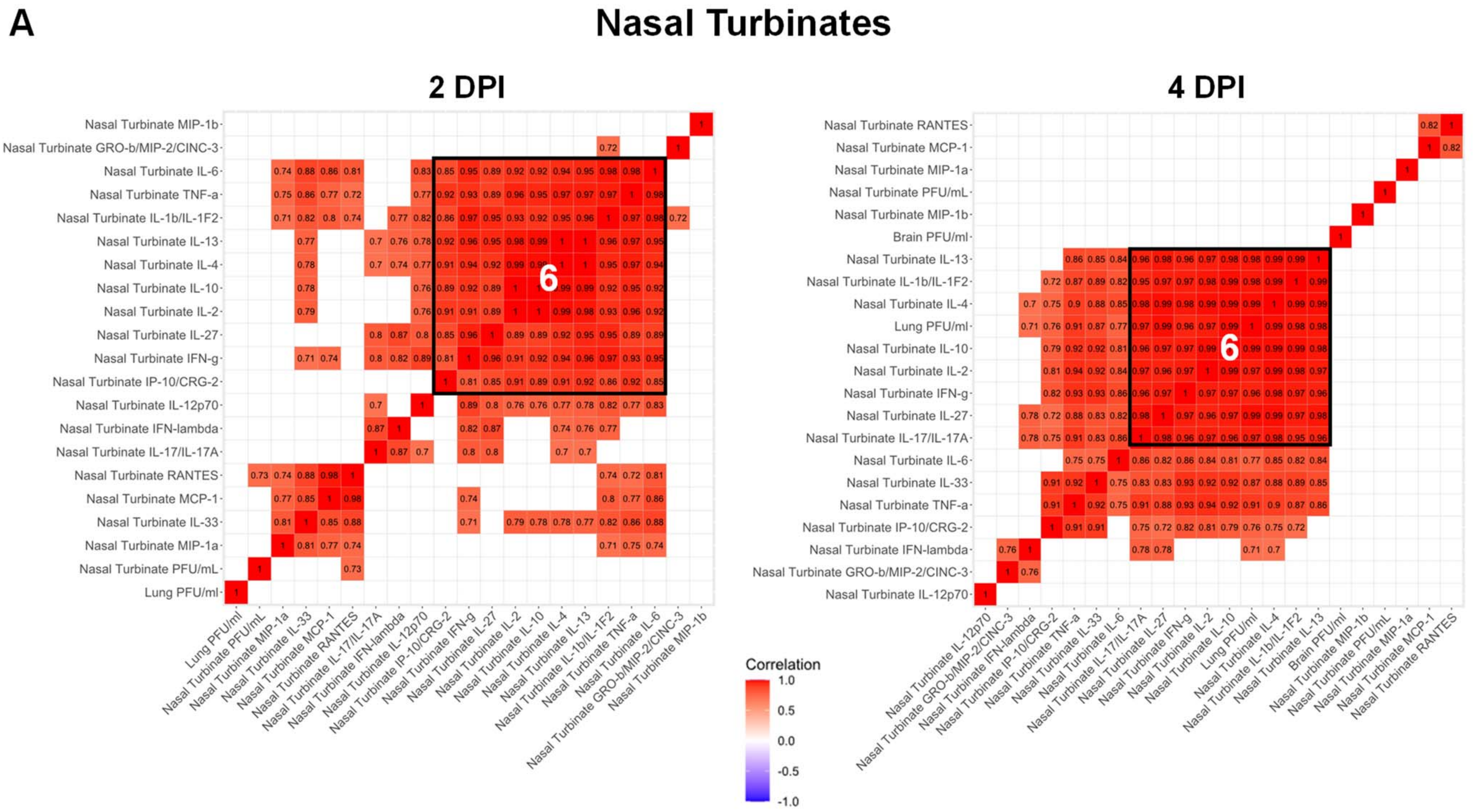


**Figure S6. SARS-CoV-2 infected K18 hACE2 transgenic mice reveal differentiated clusters of chemokine and cytokine correlations with clinical symptoms progression.** Hierarchically clustered Pearson correlations of measurements in (**A**) nasal turbinate and (**B**) trachea of SARS-CoV-2 infected K18 hACE2 transgenic mice (Left: 2-DPI, Right: 4-DPI). Positive correlation (Red = 1) and negative correlations (Blue = -1), with clusters (Black outlined boxes with cluster number). Non-significant values (p > 0.05 measured by Pearson's correlation t-test) left blank. DPI: Days post-infection. MIP-2/CXCL2; MCP-1/CCL2; MIP-1a/CCL3; MIP-1b/CCL4; RANTES/CCL5; IP-10/CXCL10.

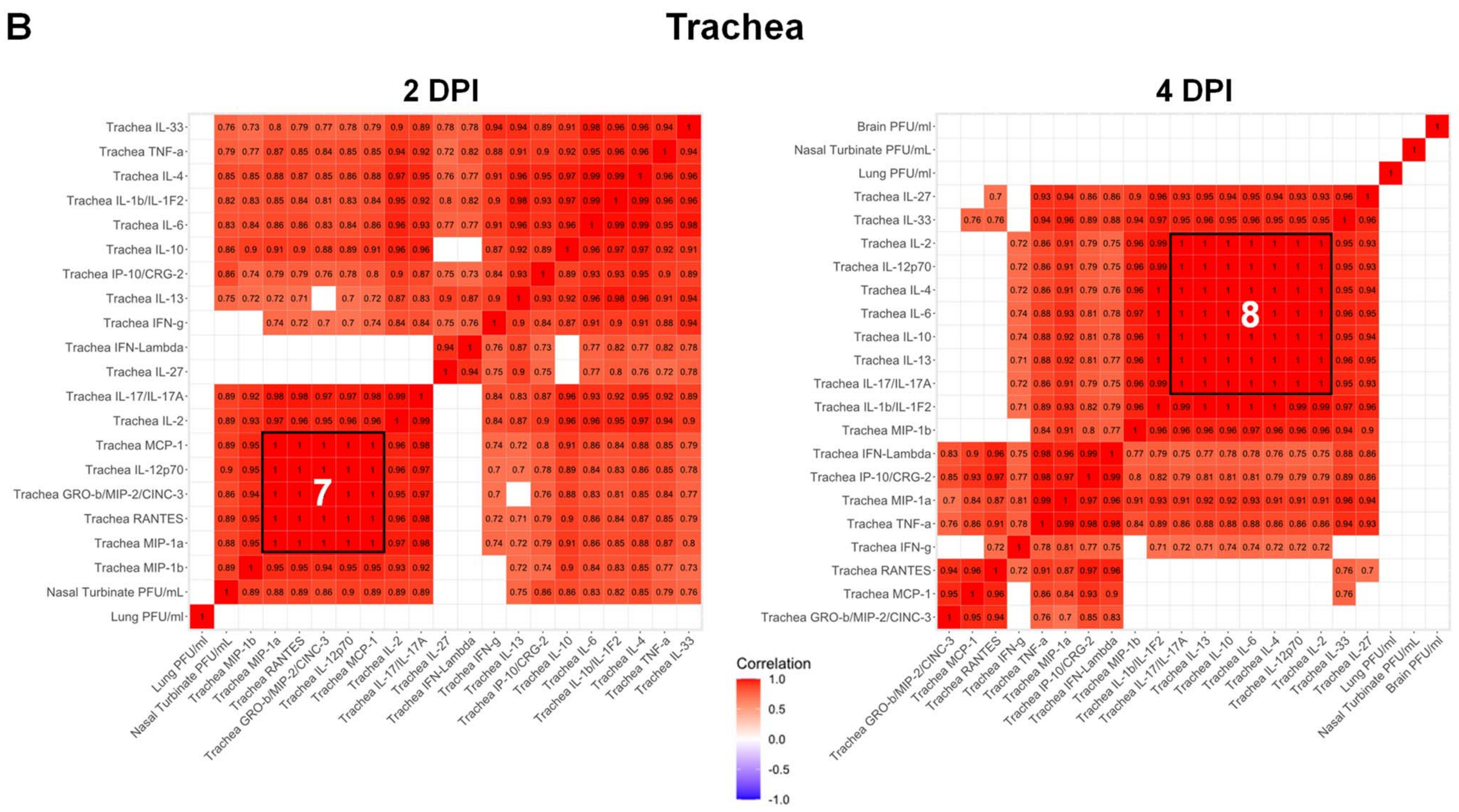


**Figure S6. Continuation.**

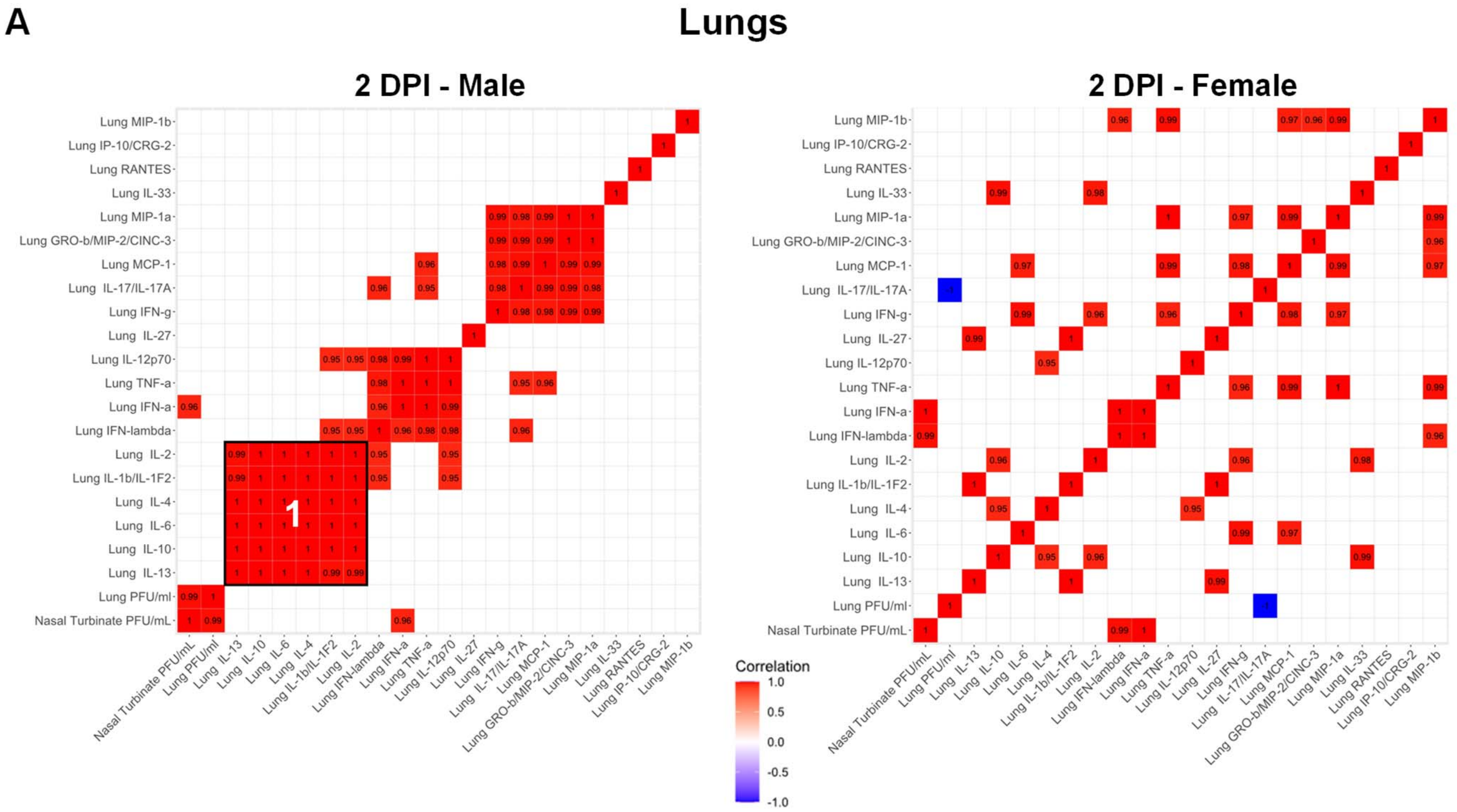


**Figure S7**. **Immune response of SARS-CoV-2 infected K18 hACE2 transgenic mice is dependent on mouse sex.** Hierarchically clustered Pearson correlations of measurements in lung at (**A**) 2- and (**B**) 4-DPI and brain at (**C**) 2- and (**D**) 4-DPI of SARS-CoV-2 infected K18 hACE2 transgenic mice (Left: Males, Right: Females). Positive correlation (Red = 1) and negative correlations (Blue = -1), with clusters (Black outlined boxes with cluster number). Non-significant values (p > 0.05 measured by Pearson's correlation t-test) left blank. DPI: Days post-infection. MIP-2/CXCL2; MCP-1/CCL2; MIP-1a/CCL3; MIP-1b/CCL4; RANTES/CCL5; IP-10/CXCL10.

**B** **Lungs**

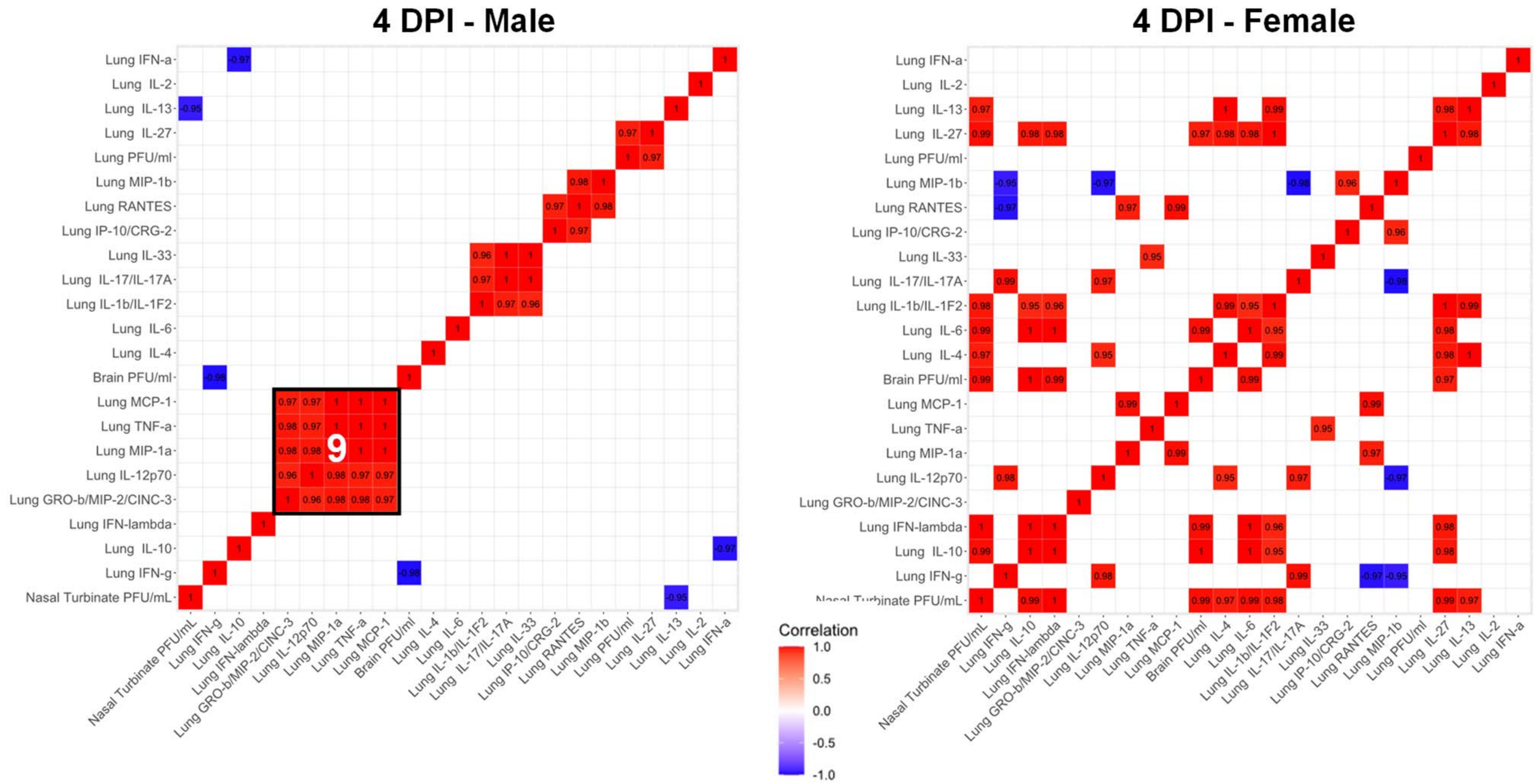


**Figure S7**. **Continuation.**

**C**

### Brain

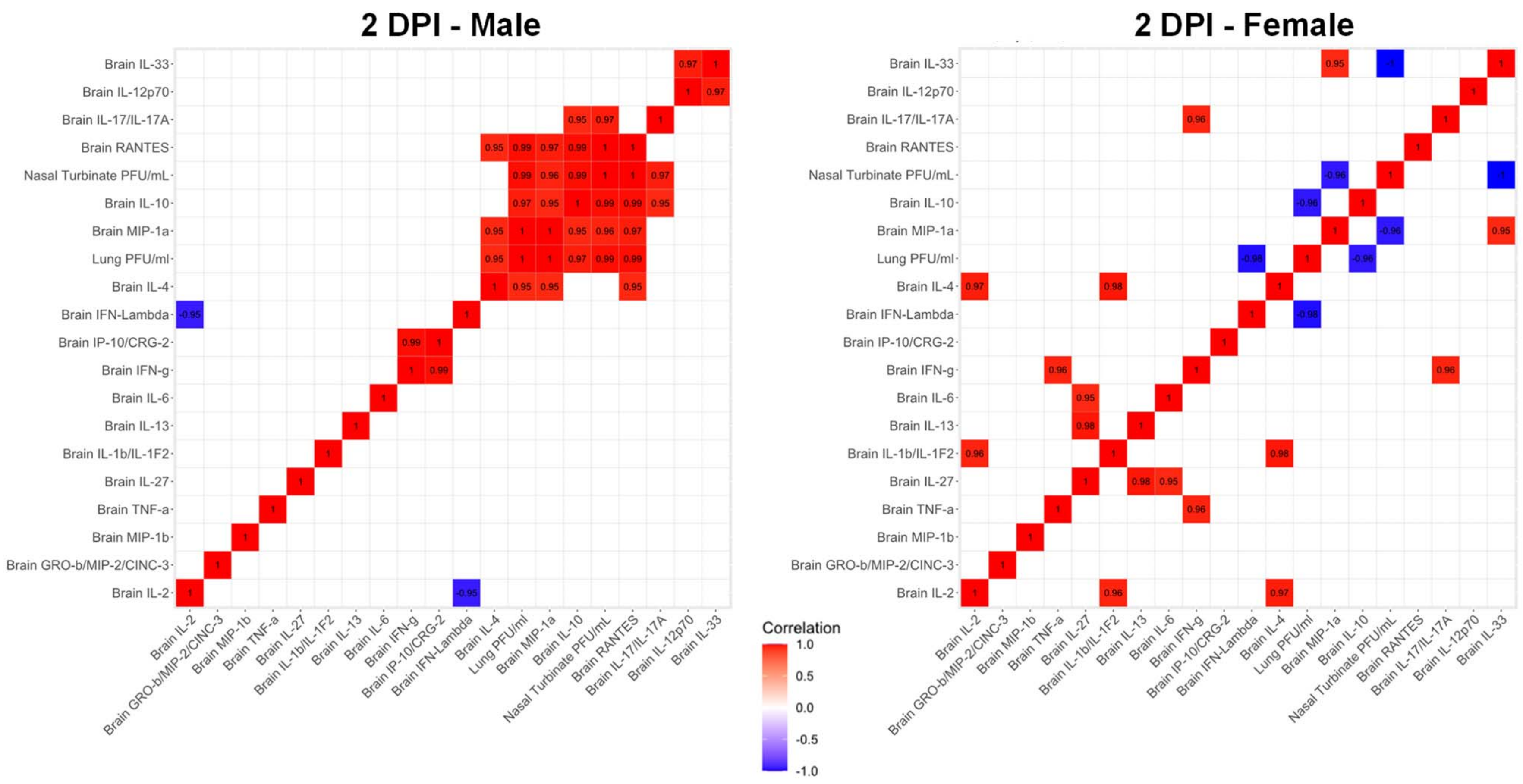


**Figure S7**. **Continuation.**

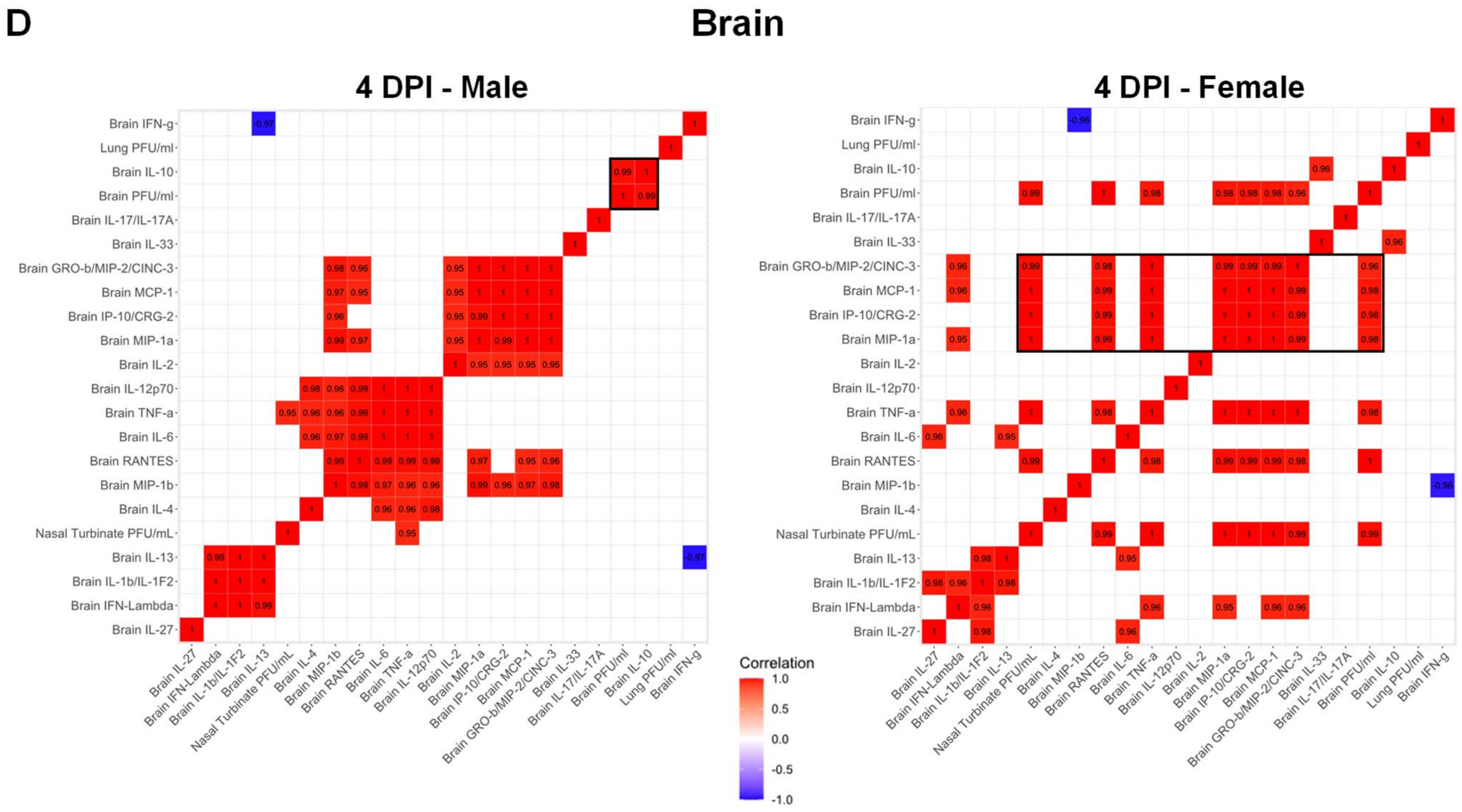


**Figure S7**. **Continuation.**
